# Two-color coincidence single-molecule pull-down for the specific detection of disease-associated protein aggregates

**DOI:** 10.1101/2023.06.27.546642

**Authors:** Rebecca S. Saleeb, Craig Leighton, Ji-Eun Lee, Judi O’Shaughnessy, Kiani Jeacock, Alexandre Chappard, Robyn Cumberland, Sarah R. Ball, Margaret Sunde, David J. Clarke, Kristin Piché, Jacob A. McPhail, Ariel Louwrier, Rachel Angers, Sonia Gandhi, Patrick Downey, Tilo Kunath, Mathew H. Horrocks

## Abstract

The misfolding and aggregation of protein is a characteristic of many neurodegenerative disorders, including Alzheimer’s and Parkinson’s disease. The wide range of sizes and structures of oligomers and fibrils generated have previously been studied using single-molecule and super-resolution microscopy. These methods, however, tend to rely on the use of either directly labeled protein, or on the addition of non-specific amyloid stains, such as thioflavin-T. This has prevented the characterization of protein aggregate composition in complex biological samples. Here, we have developed a single-molecule two-color aggregate pull-down (STAPull) assay to overcome this challenge by probing immobilized proteins using orthogonally labeled antibodies targeting the same epitope. By looking at colocalized signals, we can eliminate monomeric protein, and specifically quantify aggregated proteins. Using the aggregation-prone alpha-synuclein protein as a model, we demonstrate that this approach can specifically detect aggregates with a limit of detection of 5 pM. Furthermore, we show that STAPull can be used in a range of samples, including in human biofluids. STAPull is generally applicable to protein aggregates from a variety of disorders, and will aid in the identification of biomarkers that are crucial in the effort to diagnose these diseases.

---

Protein aggregation plays a key role in the pathogenesis of many neurodegenerative disorders, including Parkinson’s disease (PD), and Alzheimer’s disease (AD)^1^. In PD, alpha-synuclein (α-syn) is incorporated into intracellular inclusions, referred to as Lewy bodies and Lewy neurites, whereas in AD, intracellular neurofibrillary tangles of tau protein, and extracellular amyloid-beta (Aβ) plaques, are common neuropathological hallmarks. Increasing evidence suggests the early-stage oligomeric intermediates that precede the formation of these insoluble inclusions are the cytotoxic species^2,3^. Given their importance in disease pathogenesis, these oligomers are an exciting candidate biomarker of disease progression, with elevated levels of oligomeric α-syn detected in PD patient cerebrospinal fluid (CSF) by ELISA^4^. However, oligomers present as a heterogeneous population, consisting of different sizes, structures and proteoform compositions^5^, and the physical characteristics that differentiate disease-associated oligomers from benign forms are largely unknown. This is in part due to ensemble averaging approaches being unable to stratify oligomer populations. To address this, there is need for advances in technology capable of detecting and characterizing the toxic oligomeric species. Achieving this will have significant implications for patient stratification and the evaluation of clinical trial outcomes.

Protein adsorption onto glass enables direct imaging at the single-molecule level using total internal reflection fluorescence (TIRF) microscopy. This method, however, prevents the use of antibodies for labeling as they are also adsorbed onto the glass surface. We and others have successfully circumvented this hindrance by using non-specific amyloid dyes to target protein aggregates by single aggregate visualization by enhancement (SAVE) imaging^6^. Whilst SAVE and related approaches have enabled the detection^6^, and even the nanoscopic imaging of oligomers in CSF^7,8^, they rely on non-specific dyes, such as thioflavin-T (ThT), which binds a cross-beta structural motif commonly found in amyloid structures.

To overcome this and access the advantages of protein-specific immunodetection, several surface treatment methods have been developed to minimize non-specific adhesion of biomolecules to the glass surface. When it comes to single-molecule detection, single-molecule pull-down (SiMPull) is the gold standard surface passivation method^9^. In this approach, the surface is blocked by covalent modification with polyethylene glycol (PEG) molecules, a fraction of which are biotinylated to enable the specific addition of streptavidin and biotinylated antibody. The immobilized protein target is then immunolabeled and imaged using TIRF microscopy. Je *et al*. have demonstrated use of this technique to visualize total α-syn in human brain homogenates^10^, however, higher order species can only be inferred from the intensity of the fluorescent signal, resulting in an underestimation of the prevalence of oligomers in a sample.

In this work, we modified SiMPull to specifically detect protein aggregates against a background of monomeric protein using two-color coincidence detection (TCCD)^11^. Our novel approach, termed single-molecule two-color aggregate pull-down (STAPull), enables detection of picomolar levels of oligomeric α-syn, representing a 3-fold improvement in sensitivity over single-color detection. Furthermore, unlike ThT-based assays, STAPull was able to distinguish aggregates composed of different amyloid proteins with a high degree of specificity, including early-stage oligomers lacking the extended β-sheet structure required for ThT reactivity. Finally, we measured aggregates in a range of biological samples, including conditioned media and in life patient CSF, observing significantly higher titers in cases of pathology.

## Results

### STAPull distinguishes oligomeric α-syn with physiologically relevant sensitivity

In order to directly visualize α-syn oligomers, they were first immobilized on a PEGylated surface using a biotinylated anti-α-syn capture antibody tethered to the surface via biotin-streptavidin bridging^9^. Unlike SiMPull, which relies on arbitrary intensity-based filtering to delineate the oligomer population, STAPull uses a 1:1 combination of Alexa Fluor 488 (AF488)-labeled and Alexa Fluor 647 (AF647)-labeled monoclonal detection antibodies (schematized in Figure 1A). In this approach, the increased epitope availability of higher order species allows them to be separated from monomeric protein based on the colocalization of two or more orthogonally labeled antibodies. Similar dual-ELISA approaches that use the same capture and detection antibodies have been proposed^4^, however we observed that at the single-molecule level, use of a capture antibody targeting a different epitope region to the detection antibodies was critical for sensitivity (Supplementary Figure 1).

**Fig. 1.**
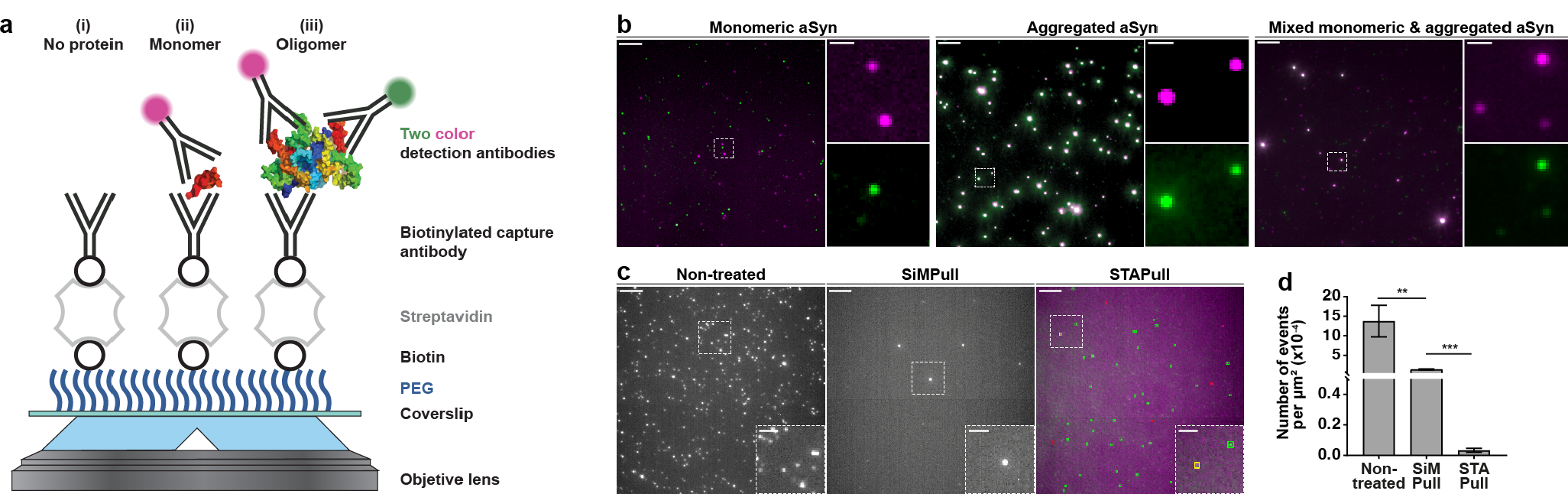
STAPull concept. **a** A glass coverslip is silanized and conjugated to PEG, of which 5% is biotinylated, and treated sequentially with streptavidin, biotinylated capture antibody and the target biomolecule to surface-immobilize. Following incubation with the sample, pulled-down protein is probed using a mixture of AF488- and AF647-labeled monoclonal detection antibody such that (i) no bound protein produces no signal, (ii) monomeric protein produces single-color signal, and (iii) oligomeric protein produces two-color signal via TIRF microscopy. **b** Demonstrative composite STAPull images for 45 nM α-syn monomer (left), 5 nM α-syn aggregates (center), or both (right). Monomer is visible as single-color puncta (green or magenta) and oligomers as two-color (white). Single-channel enlargements of the indicated regions shown. Scale bars 5 μm, crops 1 μm. **c** Representative images of the background signal resulting from adsorption of detection antibody in the absence of target protein to clean non-treated (left), SiMPull (center), or STAPull (right) surfaces. In the latter case, the channel of local maxima are indicated with colored boxes to highlight whether single channel (green and red) or two-channel (yellow). Scale bars 5 μm, insets 2 μm. **d** Mean density and standard deviation of detections in c (n = 3, 16 technical repeats), statistical significance determined with an unpaired student t-test, **p < 0.01, ***p < 0.001.

As recombinant α-syn can self-assemble into fibrils, accelerated by agitated incubation at 37 °C, we prepared monomeric and aggregated samples as an *in vitro* model of pathology. Application of nanomolar concentrations of monomeric or aggregated protein to the STAPull surface demonstrated increased two-color coincident events in the latter case, indicative of oligomeric species (Figure 1B). Based on the ratio of coincident to total detections, we observed that aggregated α-syn was composed of ∼44.5% oligomeric species, while ‘pure’ monomer contained ∼6.8% oligomer. Combining these two solutions at a molar ratio of 1:9 produced a solution containing 11.6% aggregated protein as would be theoretically anticipated based on the above stoichiometry.

The primary aim of surface passivation is to block non-specific protein adsorption, however perfect passivation is unfeasible and low-level background signal persists^12^. STAPull minimizes this residual background by rejecting single-channel non-specific events alongside monomeric signal. To demonstrate this, we compared the background signal produced by exposing SiMPull, STAPull or non-treated surfaces to 1 nM labeled antibody in the absence of target protein (Figure 1C-D). SiMPull surface passivation provides a ∼10-fold reduction in non-specific binding compared with a non-treated surface (mean ± SD: 13.77×10^−4^ ± 4.03×10^−4^ events/μm^2^ vs. 1.40×10^−4^ ± 0.12×10^−4^ events/μm^2^). This can be enhanced by a factor of ∼40 when only coincident events are measured using STAPull (0.03×10^−4^ ± 0.01×10^−4^ events/μm^2^) (Figure 1D).

To determine whether this reduction in background signal led to improved detection sensitivity, we measured a dilution series of aggregated α-syn and determined the lowest concentration of protein that could be reliably detected^13^. STAPull produced a limit of detection (LoD) of 22.6×10^−4^ events/μm^2^, or 358 pM monomer equivalent α-syn, which was a 3-fold improvement in sensitivity compared to SiMPull and near 600-fold improvement on ThT fluorimetry (Figure 2A-C). To approximate the oligomer concentration this equates to, we used the average intensity of single-channel events in a pure monomer sample to estimate the average number of monomer units per aggregate as 30.5 monomeric units, which aligns well with previously validated oligomer sizes^14^. When accounting for both oligomer size and the 44.5% oligomer load determined above, STAPull has an approximate LoD of 5 pM oligomer concentration. Importantly, this brings STAPull sensitivity into a physiological range, with the oligomer content of PD patient CSF determined by ELISA to be ∼10 pM^15,16^. We attribute this improvement over intensity-thresholded SiMPull to both STAPull’s increased signal-to-background ratio and its independence from sample or technique-derived intensity inconsistencies.

**Fig. 2.**
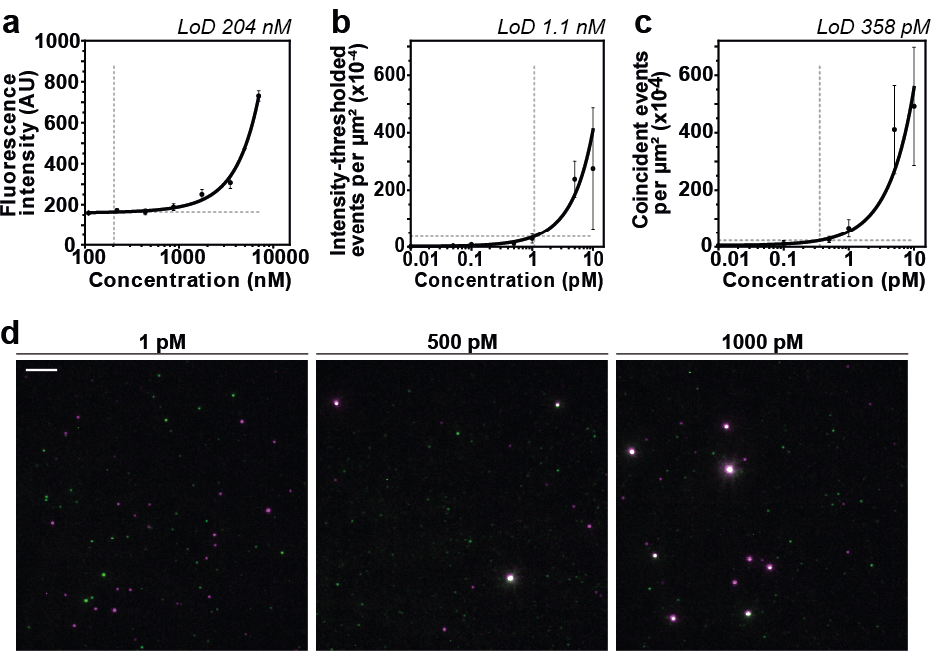
Improved limit of detection using STAPull. **a** ThT fluorescence spectroscopy of α-syn aggregates as a function of monomer-equivalent α-syn concentration. **b** Single-molecule α-syn aggregate density measured by intensity-thresholded SiMPull or **c** STAPull as a function of monomer-equivalent α-syn concentration. Dashed lines indicate the limit of detection (LoD) and the corresponding concentration is determined via a two-order polynomial fit (black line). Data is mean of n = 3 (each averaged over 64 technical repeats) and standard deviation shown. **d** Representative images of STAPull with α-syn aggregate concentration below (1 pM), near (500 pM) and above (1000 pM) its LoD. Scale bar 5 μm.

### Specific detection of protein aggregates using STAPull

SAVE imaging can achieve similar detection sensitivity to STAPull, but its application is limited by an inability to determine protein identity and thus disease type. Conversely, as an epitope-mediated assay, STAPull has the capacity to differentiate the underlying proteinopathy. To verify this, we prepared three different surface types, (i) a poly-L-lysine coating to facilitate total protein adsorption for SAVE-based ThT detection, (ii) a STAPull surface with α-syn specific capture and detection antibodies, (iii) a STAPull surface with tau specific capture and detection antibodies. Upon exposure of each surface to aggregates of either amyloid-beta (Aβ), α-syn, tau, or buffer alone, only the STAPull surfaces exclusively detect the target protein for which they were designed (Figure 3A). ThT detection, on the other hand, demonstrates non-specific signal elevation compared to the protein-free control in response to all three amyloid proteins. Notably, the densities of α-syn and tau observed on PLL (1344.95×10^−4^ and 14.58×10^−4^ events/μm^2^, respectively) are broadly similar, or elevated, on STAPull surfaces (500.72×10^−4^ and 195.58×10^−4^ events/μm^2^), despite applying a 50-fold lower concentration to the latter. This highlights the importance of the capture antibody for protein enrichment on the surface and the suitability of STAPull for probing rare oligomeric species in patient biofluids.

**Fig. 3.**
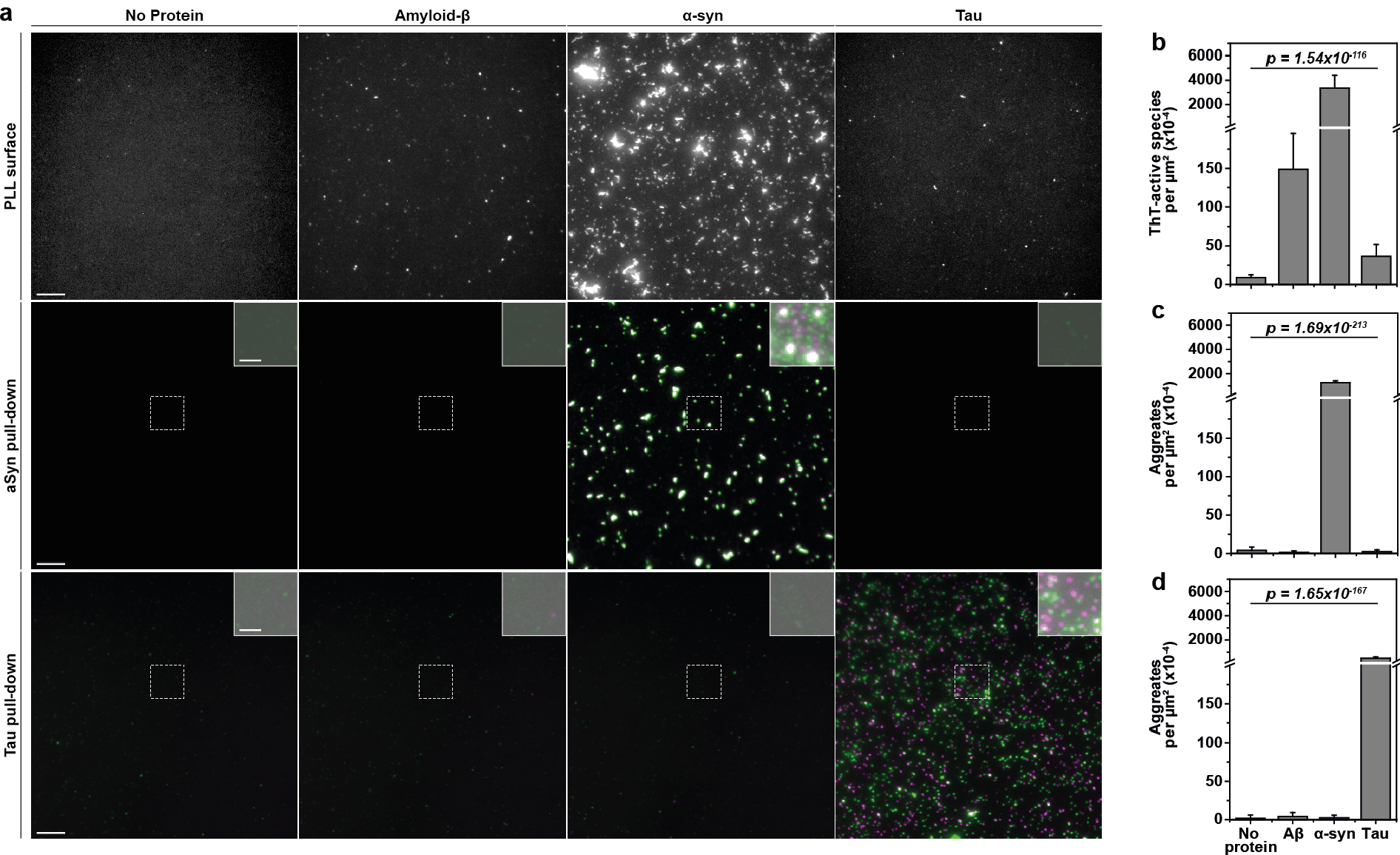
STAPull provides protein-specific aggregate detection. **a** Representative TIRF microscopy images of different surface preparations exposed to aggregated Aβ, α-syn, tau or no protein as indicated. Protein probed with (top row) ThT by SAVE imaging on a PLL-coated surface, (middle row) two-color antibodies against α-syn on an α-syn-specific STAPull surface, or (bottom row) two-color antibodies against tau on a tau-specific STAPull surface. Enlarged insets of indicated regions are at high contrast with 0.5 gamma adjustment to visualize any low-level signal. Scale bars 5 μm, inset 2 μm. **b** Mean density of ThT-reactive species detected on the PLL surface, **c** mean density of two-color species on the α-syn STAPull surface and **d** the tau surface. A split y-axis is used to visualise low-level variation (scaled 0-200 and 250-7000 counts per μm^2^ (×10^−4^) pre- and post-break, respectively). Standard deviation of 64 technical repeats shown with statistical significance determined by one-way ANOVA analysis (refer to Supplementary Table 1 for details of post hoc Tukey pairwise means comparisons).

Though it would be plausible to create specificity for ThT detection via the capture antibody alone, we and others^12^ have observed that PEG passivation is unable to completely resist protein adsorption to the surface, necessitating antibody detection to achieve specificity (Supplementary Figure 2). Furthermore, whereas STAPull can theoretically detect dimeric species, ThT is only able to bind amyloid structures containing extended β-sheet structure^17^.

### STAPull detection of kinetically stable oligomers

We observed that upon assaying different timepoints of α-syn exposed to conditions favoring its aggregation, it was possible to identify oligomers with STAPull earlier than with ThT detection (Supplementary Figure 3). As early oligomeric structures are increasingly believed to be the neurotoxic species driving synucleinopathy^2^, and are most likely present at the earliest stages of disease, a means to detect them would have significant implications for early diagnosis, drug discovery and the evaluation of clinical trial outcomes. We therefore sought to assess the suitability of STAPull for the detection of these species using kinetically stable oligomers.

Kinetically stable oligomers have previously been characterized as soluble globular α-syn structures with toxic properties demonstrated *in vitro*^18^. They appear to have a β-sheet structure intermediate between that of the monomer and the fully structured amyloid fibril^18^. We used two-color coincidence detection and a ThT post-stain to concurrently assay commercially obtained α-syn monomer, kinetically stable oligomers, and pre-formed fibrils using both STAPull and SAVE imaging. The aggregate species were structurally validated by transmission electron microscopy (Supplementary Figure 4) to confirm globular and fibrillar structure, respectively. With SAVE imaging, we observed a significant elevation in ThT puncta for the fibrillar species as compared to monomer (mean per μm^2^ ± SD: 312.38×10^−4^ ± 177.80×10^−4^and 79.24×10^−4^ ± 98.21×10^−4^, respectively), and these correlated with coincident STAPull detections (Figure 4A-B). However, no increase was evident for the oligomer sample (13.57×10^−4^ ± 10.46×10^−4^), indicating a population of toxic oligomeric species that cannot be detected by ThT-based assays. In contrast, STAPull demonstrated a significantly increased prevalence of coincident detections for both oligomers and fibrils (mean per μm^2^ ± SD: 2055.54×10^−4^ ± 123.44×10^−4^ and 775.37×10^−4^ ± 83.30×10^−4^ respectively) as compared to monomer (160.58×10^−4^ ± 90.53×10^−4^), confirming the ability of STAPull to detect these early oligomeric species (Figure 4C).

**Fig. 4.**
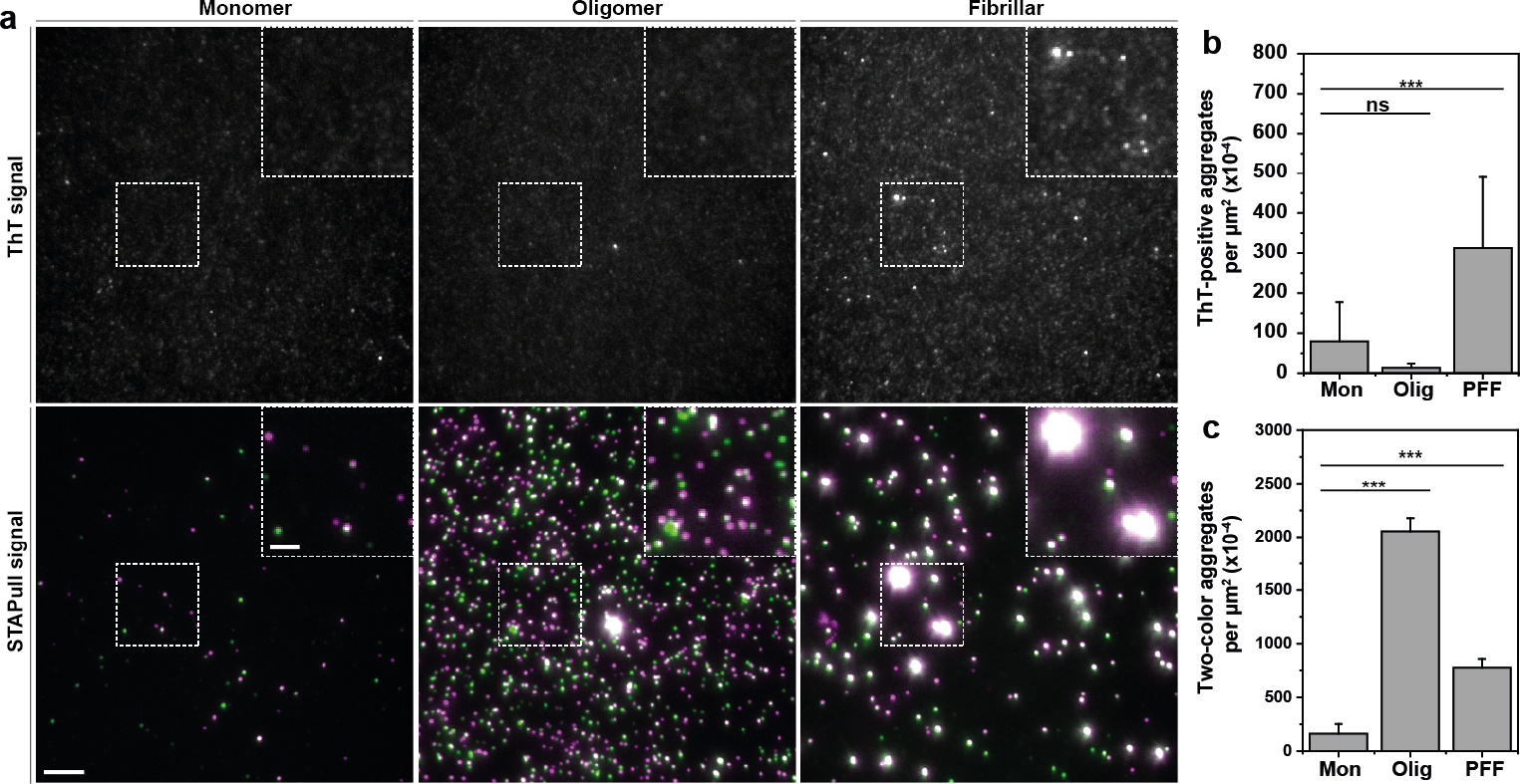
STAPull detects ThT-negative α-syn oligomers. **a** Representative single-molecule pull-down images of commercially sourced human recombinant α-syn in the monomeric, kinetically stable oligomeric or pre-formed fibrillar form as indicated. The same field of view is presented with ThT detection (top) or STAPull two-color coincidence (bottom), with the boxed region magnified in inset. Scale bar 5 μm in full-field and 2 μm in inset. **b** Quantification of the mean number of ThT puncta or **c** STAPull coincident events per μm^2^ for the dataset represented in a. Error bars indicate standard deviation of 25 technical repeats, statistical significance was determined with an unpaired one-tailed student’s t-test, ***p < 0.001.

### STAPull detection of endogenous protein aggregates

The data presented thus far has demonstrated the suitability of STAPull for the specific detection of early α-syn oligomers generated *in vitro*. Evidence suggests these species are driving neurotoxicity^2^, but their detection in biological samples is made challenging due to the complex milieu of biomolecules that may impede immunocapture, non-specifically bind antibodies or elevate baseline signal through autofluorescence. Therefore, to determine the suitability of the technique for probing α-syn in such samples, we first used culture medium exposed to human midbrain dopaminergic (mDA) neurons, the primary cell type lost in PD. This ‘conditioned media’ acquires secreted biomolecules and cellular cargo from the cultured cells, thus resembling extracellular fluid and offering a useful *in vitro* model.

mDA neurons were differentiated from induced pluripotent stem cells (iPSCs) derived from a PD patient harbouring a triplication of the α-syn-encoding *SNCA* gene (*SNCA*^*Trip/+*^) or its CRISPR-engineered knockout (*SNCA*^*-/-*^). This mutation is known to cause autosomal dominant, early-onset PD due to increased α-syn burden^19,20^ and iPSC derived neurons exhibit elevated levels of α-syn aggregates^21^. By using STAPull to assay 3-day conditioned media, we observed a significantly higher total α-syn titre in the *SNCA*^*Trip/+*^ medium than that of medium from *SNCA*^*-/-*^ neurons (mean per μm^2^ ± SD: 2528×10^−4^ ± 367×10^−4^ and 762×10^−4^ ± 170×10^−4^, respectively). Indeed, the knockout cells showed no significant elevation above the baseline detections observed in cell-free medium (692×10^−4^ ± 66×10^−4^) (Figure 5A-B). In addition, the number of α-syn aggregates present, indicated by the number of events with coincident signal, also trended upwards in the case of *SNCA*^*Trip/+*^ alone (Figure 5C), demonstrating that STAPull is sensitive to differences in α-syn oligomer burden in a complex sample.

**Fig. 5.**
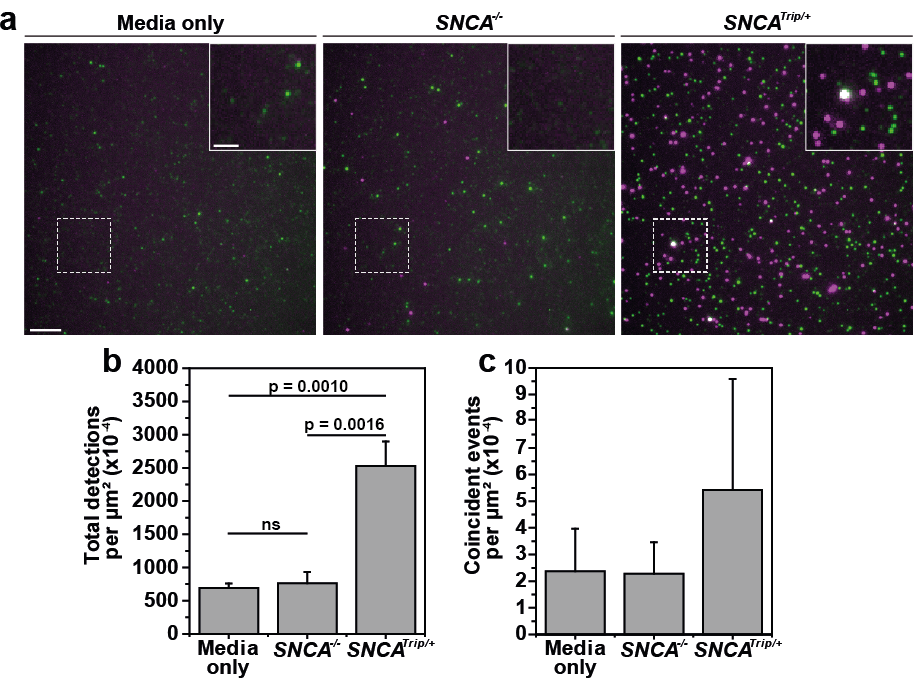
STAPull detects elevated α-syn in iPSC-derived mDA-conditioned media. **a** Representative STAPull images of 3-day conditioned culture medium in the absence of cells (left), and in the presence of human iPSC-derived mDA neurons that harbour either a *SNCA* knockout (*SNCA*^*-/-*^, center) or triplication of the *SNCA* locus (*SNCA*^*Trip/+*^, right). Monomeric α-syn appear as single-color puncta (green or magenta), and oligomeric appears as two-color puncta (white). Region indicated is magnified in inset. Scale bar 5 μm, inset 2 μm. **b** Mean density of all detected species (monomeric and oligomeric) and **c** coincident only species, for dataset presented in a. Error bars show standard deviation (n = 3, each with 64 technical repeats) and statistical significance determined with an unpaired student t-test.

Given the sensitivity of STAPull for the detection of secreted α-syn aggregates, we next sought to assess the approach for the detection of aggregates in patient *ante-mortem* CSF, which had been previously characterised by ELISA^22^ and SAVE imaging^6^ (demographics, clinical features and biomarkers outlined in Supplementary Table 2). As STAPull is effective in minimizing background, 50 μL CSF could be applied without dilution to maximize α-syn enrichment on the surface. CSF from PD patients (N = 7), AD patients (N = 5) or healthy individuals (N = 13) were probed with an antibody targeting the N-terminal region of α-syn to ensure detection of C-terminally truncated protein commonly associated with PD pathology^23^. The data showed that, akin to published SAVE data^6^, a significant increase in the α-syn aggregate number is observed for individuals diagnosed with PD (8.05 ± 1.43 fold above PBS baseline aggregate number), as compared to healthy controls (4.44 ± 1.28 fold) giving an ANOVA p-value of 7.47×10^−6^ (Figure 6). Importantly, STAPull did not detect a significant elevation in α-syn aggregates in the AD patient CSF (mean ± SD: 4.98 ± 0.57, p-value 0.68) versus healthy control, and thus, unlike SAVE imaging, is able to specifically differentiate the elevated α-syn burden in PD patient biofluids (Figure 6).

**Fig. 6.**
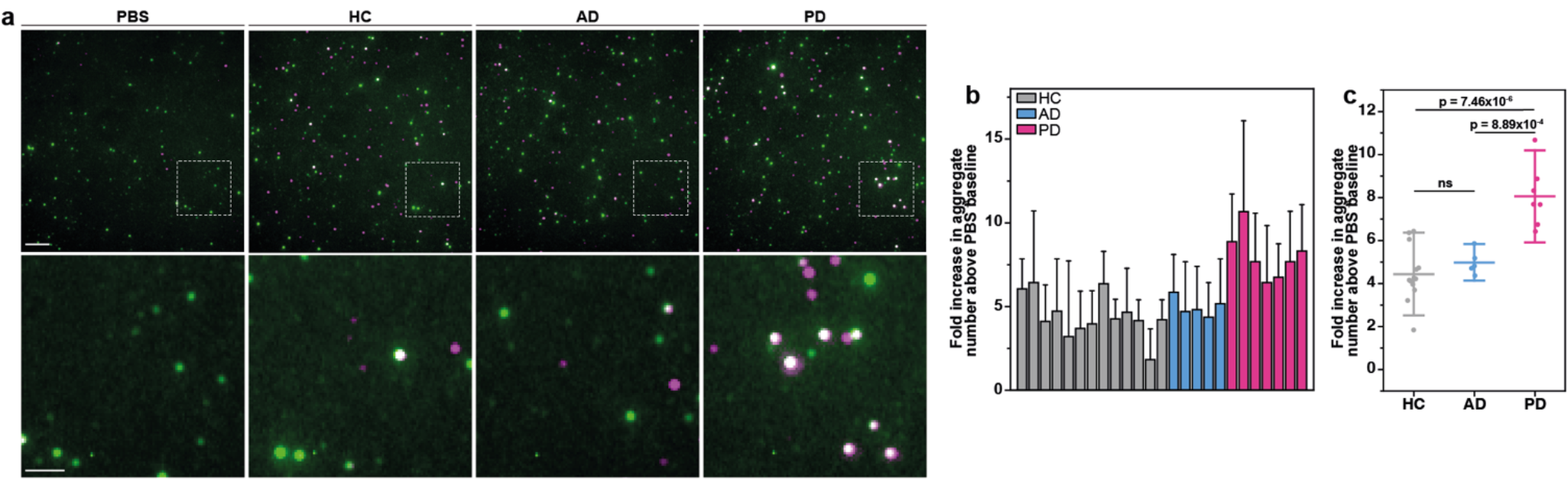
STAPull detects α-syn aggregates in PD patient CSF. **a** Representative STAPull images of in life CSF from patients diagnosed with PD, AD or healthy control individuals (HC) and in comparison to a PBS-only control (as indicated). The boxed region indicated is enlarged in the row below. Monomeric species are visible as single-color puncta (green or magenta) and oligomeric as two-color (white). Scale bars 5 μm (top), 2 μm (bottom). **b** Quantification by case of aggregate density given as fold-increase above PBS baseline, error bars indicate standard deviation of 64 technical repeats. **c** Dot plot comparing the data in b by diagnosis, where mean is indicated by long horizontal line and standard deviation by vertical error bars. Statistical significance between diagnostic groups was confirmed by one-way ANOVA (p-value 9.88×10^−6^), where PD samples alone differed from other groups based on Tukey means comparison (p-values indicated).

## Discussion

At present, neurodegenerative diseases are clinically diagnosed by the manifestation of symptoms that follow irreversible neuronal cell death. However, the neurodegenerative process begins 10-20 years prior to symptom onset, and such late diagnosis precludes the introduction of disease-modifying therapeutics when they could have the greatest effect. Indeed, this issue is considered a major contributor to the failure of late-stage clinical trials^24^ and underlines the need for biomarker-based diagnostic tools to detect neurodegeneration at its earliest stages.

Ensemble immunoassays such as ELISA or SiMoA are the gold standard approach for protein biomarker detection, and their adaptation for aggregated protein pull-down have demonstrated oligomer-specific detection^4,15,16^. However, these approaches detect oligomeric α-syn in the CSF of both healthy (accounting for ∼4% of total α-syn) and PD patients (∼9% of total α-syn), highlighting the need to dissect pathological from benign species, particularly in early-stage patients where elevation alone may be incremental. Techniques such as SAVE and SiMPull have attempted to address the need for aggregate species stratification through single-molecule visualization. Indeed, SAVE imaging has observed different aggregate subpopulations in healthy versus disease cases based on SAVE intensity^6^, but cannot specify protein identity. SiMPull is able to build on this by offering protein-specificity alongside single-molecule detection, however it lacks the sensitivity for diagnostic applications due to high background from monomeric protein and non-specific binding.

We describe here a novel approach, STAPull, to address these issues. STAPull enables single-molecule detection and differentiation of protein-specific oligomers, the earliest stage aggregate species associated with neurotoxicity. We use detection of α-syn, the hallmark aggregating protein of synucleinopathies such as PD, to demonstrate picomolar sensitivity capable of detecting significantly higher oligomer burdens in PD patient CSF as compared to healthy control cases. This approach is broadly applicable to the detection of any proteinopathy or multimeric structures by replacement of the capture and detection antibodies. Furthermore, STAPull does not require the use of oligomer-specific antibodies that may only detect specific conformations, but instead can be used with antibodies targeting the constituent protein.

The key advantage of STAPull is its ability to identify and visualize protein aggregates, providing opportunity for the discrimination of aggregate subspecies. As different oligomer strains have been identified in a number of neurodegenerative diseases, including PD, AD and CJD^25–27^, the detection of subpopulations with different pathological properties may provide a powerful prognostic biomarker. Towards this end, incorporation of super-resolution imaging techniques, such as DNA-PAINT or dSTORM, could enable oligomer stratification based on morphological metrics.

In summary, we present a novel technique that enables, to our knowledge, the first direct detection of specific protein aggregates at the single-molecule level. This tool has immediate implications for aiding the evaluation of aggregation-inhibiting drugs and clinical trial outcomes, and may present opportunities for early-stage diagnostics.

## Methods

### Preparation of α-syn

Wild-type α-syn (Addgene, 36046^28^) was expressed and harvested from *Escherichia coli* as described previously^29^. The cell pellet was resuspended in 20 mM Tris-HCl pH 8.0, 1 mM EDTA (buffer A) supplemented with 1 mM PMSF, lysed by 15 min pulsed probe sonication and clarified by 12,000 RCF centrifugation at 4 °C. The supernatant was recovered, heated to 80 °C for 10 min and centrifuged at 12,000 RCF. 10 mg/ml streptomycin sulphate (086K1263) was added to the supernatant and incubated 20 min at 10 °C with 100 rpm shaking. Following centrifugation at 12,000 RCF, the supernatant was collected and the previous step repeated with 360 mg/ml ammonium sulphate (101575102). The pellet was collected by 45 min centrifugation at 12,000 RCF, resuspended in 25 mL buffer A and dialysed overnight at 4 °C in buffer A. The sample was loaded onto a HiTrap Q FF column (GE Healthcare Life Sciences) on an ÄKTA Start Protein Purification System (GE Healthcare Life Sciences). α-Syn was eluted at approximately 300 mM NaCl with a salt gradient from 0 mM to 1000 mM. The eluent was then loaded into a HiPrep 26/60 Sephacryl S-200 size exclusion column and eluted in 20 mM Tris-HCl pH 7.4, 100 nM NaCl. Fractions corresponding to the UV chromatograph peak were recovered and confirmed by SDS-PAGE, then concentrated using a 5 kDa spin column (Sartorius, VSO611) and flash-frozen with liquid nitrogen in single-use aliquots for storage at -80 °C. The protein purity was quality checked by LC-MS on a quadrupole ion-mobility time of flight instrument (Synapt G2, Waters Corp.).

### *In vitro* aggregation of α-syn

Monomeric wild-type α-syn was ultracentrifuged at 90,000 RCF for 1 hour at 4 °C to remove any amorphous aggregates (Beckman Coulter Optima Max-XP with MLA-130 rotor). The supernatant was recovered and the concentration determined by absorbance at 275 nm using an extinction coefficient of 5,600 M^-1^ cm^-1^. The concentration was adjusted to 70 μM by dilution in aggregation buffer (0.02 μm-filtered 20 mM Tris-HCl pH 7.4, 100 nM NaCl, 0.01% (w/v) NaN_3_). The protein was then incubated at 37 °C for 120 hours with constant agitation at 200 rpm and the product sonicated 10 min at 10 °C (0.5 min on/0.5 min off) using a Biorupter Pico (Diagenode). Aggregates were flash-frozen with liquid nitrogen in single-use aliquots for storage at -80 °C.

### Generation of α-syn kinetically stable oligomers and pre-formed fibrils

Human α-syn monomer, oligomers and pre-formed fibrils are available from a commercial source (StressMarq Biosciences Cat# SPR-321, SPR-484 and SPR-322, respectively), which are generated using methods previously described. Prior to oligomerization/fibrilization, α-syn monomer was thawed, mixed, and centrifuged 20,000 xg at 4 °C to remove any pre-existing aggregate. The supernatant containing monomer was pooled and re-filter sterilized (0.2 µm). To generate kinetically stable oligomers, monomeric α-syn was diluted to 2 mg/mL in 1x phosphate buffer pH 7.4, evaporated in a Speedvac (Savant) and re-suspended in dH_2_O a total of fifty cycles. Oligomers were pooled, washed on a 100 kDa MWCO concentrator (Amicon), concentrated to 2 mg/mL, filter sterilized (0.2 µm), aliquoted and frozen to -80 °C. Oligomers were confirmed by native PAGE, size-exclusion chromatography and transmission electron microscopy (TEM). To generate pre-formed fibrils (PFFs), monomers were diluted to 5 mg/mL with PBS pH 7.4 and shaken at 1000 rpm, 37 °C, for seven days using a Thermomixer C with a heated lid. PFFs were aliquoted, frozen at -80 °C, and confirmed by sedimentation assay, ThT response and TEM.

### Aggregation of Aβ

Commercial Aβ 1-42 monomer (Anaspec, AS-20276) was prepared at 2 μM in saline sodium phosphate EDTA (SSPE) buffer and aggregated by 48 hour incubation at 37 °C. Single-use aliquots were flash-frozen in liquid nitrogen and stored at -80 °C.

### Aggregation of tau

Tau4R monomer was diluted to 20 μM in SSPE buffer supplemented with 2 μM heparin and 0.01% (w/v) NaN_3_. Protein aggregation was carried out by 14-day incubation at 37 °C, following which aliquots were flash-frozen in liquid nitrogen and stored at -80 °C.

### Thioflavin-T preparation

ThT stock solutions were prepared by diluting ThT (Abcam, ab120751) into neat ethanol to give a final concentration ∼5 M. The solution was vortexed thoroughly, and a working stock prepared at ∼200 μM by dilution in PBS. The solution was filtered through a 0.02 μm syringe filter (Whatman, FIL2824) to remove insoluble ThT that may give rise to fluorescent puncta and the concentration was confirmed by absorbance at 412 nm, using an extinction coefficient of 36,000 M^-1^ cm^-1^. The working stock was stored in the dark at 4°C and used for a maximum of 4 days with fresh filtration on day of use

### Antibody labeling

All antibodies were generated by UCB Biopharma using single B cell technology and selected based on affinity to human fibrillar α-syn as determined by surface plasmon resonance. Details of all antibodies are outlined in supplementary table 3. Stable antibody conjugation was carried out by reaction of the antibody primary amino groups with either NHS ester-linked AF488/AF647 dyes or NHS-PEG_4_-biotin for the detection and capture antibodies respectively. For this purpose, commercial labeling kits were used (A20181/6, ThermoFisher Scientific and Pierce, 90407, respectively) in accordance with the manufacturer’s instructions. The concentration of the resultant conjugate was determined based on absorbance at 280 nm, using an extinction coefficient of 210,000 M^-1^ cm^-1^, corrected in the case of the dye-labeled conjugates using absorbance at 494 nm or 650 nm as applicable as per the manufacturer’s instructions.

### STAPull surface preparation

24 × 60 mm borosilicate glass coverslips (VWR, 631-1339) were exposed to argon plasma for 45 min to remove organic contaminants. They were subsequently submerged in 0.22 μm-filtered 1 M KOH for 20 min to activate the surface for silane functionalisation, rinsed in deionised water and transferred to 1% (3-aminopropyl)trimethoxysilane (Sigma-Aldrich, 281778) in methanol supplemented with 5% acetic acid for 20 min incubation in the dark. Following silanization, the coverslips were serially washed in methanol and deionised water, blast-dried with argon gas and affixed to an 18-well gasket (Ibidi, 81818). 50 μL freshly prepared PEG solution (100 mg/mL mPEG-SVA (5000 Da, Laysan Bio), 5 mg/mL biotin-mPEG-SVA (5000 Da, Laysan Bio) and 0.1 M NaHCO_3_) was applied to each well and incubated overnight. The surface was then rinsed with deionised water and blast-dried with argon gas, before incubating 10 min with 0.2 mg/mL streptavidin (ThermoFisher Scientific, 21125) in 0.02 μm-filtered T50 buffer (10 mM Tris-HCl pH 8.0 supplemented with 50 mM NaCl) and subsequently washed three times in T50.

To specifically enrich the target protein on the surface, 100 nM of appropriate capture antibody diluted in 0.02 μm-filtered PBS was applied for 20 min incubation (SYN-CT1-biot for α-syn and anti-tau-biot for tau pull-down). Excess antibody was removed by washing three times with PBS and 50 μL sample applied. Unless otherwise stated, recombinant protein generated in-house was applied at 10 nM concentration for 20 min, commercially sourced α-syn monomer, oligomer and fibrils (Stressmarq, SPR-321, SPR-484 and SPR-322) were applied at 25 nM concentration supplemented with 1% bovine serum albumin, conditioned media was applied neat for 20 min and CSF applied neat for 24 hours at RT. Wells were washed three times with PBS to remove unbound protein and incubated with a 1:1 mixture of 1 nM AF488- and 1 nM AF647-labeled detection antibody diluted in T50 for Tris-mediated quenching of any remaining unbound dye. Recombinant protein and conditioned media were incubated for 10 min at RT with SYN-CT2 or anti-tau for α-syn and tau detection, respectively. To ensure detection of C-terminally truncated α-syn aggregates commonly associated with PD, we alternatively used a lower affinity N-terminal-targeting SYN-NT1 antibody to probe α-syn aggregates in human CSF, applying a longer 24 hour incubation at RT. Excess detection antibody was removed by washing three times with PBS.

### Surface preparation for SAVE imaging

Coverslip surfaces were prepared as described previously^6^. Briefly, coverslips were exposed to argon plasma for 45 min to remove auto-fluorescent organic contaminants. Frame-seal slide chambers (Biorad, SLF-0601) were affixed to the glass, and 50 μL of poly-L-lysine (PLL, Sigma-Aldrich, P4707-50ML) was incubated in the chamber for 15 min. Excess PLL was removed by washing three times with 0.02 μm-filtered PBS. 500 nM protein was incubated on the surface for 1 hour and replaced with 5 μM ThT solution immediately prior to imaging.

### Single-molecule imaging

All samples were imaged using an ONI Nanoimager equipped with a 100x/1.4 NA oil immersion objective lens and ORCA-Flash 4.0 V3 sCMOS camera. Samples were exposed sequentially to 638 nm, 488 nm and/or 405 nm excitation by total internal reflection using a 53.5 degree illumination angle for visualization of AF647, AF488 and ThT as required. The resultant emission was split with a 640 nm dichroic, projecting emission above and below this wavelength onto different regions of the camera chip for dual-channel viewing. Each field of view (FOV) was imaged for 10 frames at a rate of 20 frames second^-1^ and an 8 × 8 grid of 200 μm-spaced FOVs were captured per condition to account for any region-specific variations.

### Image pre-processing

Single channel images were cropped from the dual-view acquisition and registered based on ground-truth TetraSpeck microsphere (ThermoFisher Scientific, T7279) data captured alongside the dataset. The imreg_dft Python library^30^ was used to perform and automate channel registration. Single channel image series were subsequently maximum intensity projected to reduce image noise.

### STAPull two-color coincidence analysis

Particle detection, colocalization, and quantification were performed using the ComDet v.0.5.5 plugin for ImageJ (https://github.com/ekatrukha/ComDet). To exclude background fluorescence variations, detections smaller than the resolution limit of the technique were excluded and an intensity threshold set to 5 SD above image mean was used. This value was determined empirically by assessing the number of events detected in the presence and absence of detection antibody (supplementary figure 5). Two-color events with a centre of mass within two-pixel proximity were considered coincident. To assess chance coincidence due to sample density, the same measurements were applied to a transformed control of each FOV, whereby one channel was flipped horizontally and vertically with respect to the other. Presented aggregate values represent the mean coincidence of all FOVs less their chance detections, calibrated to the imaged area. Where reported, total protein values represent the sum of single channel events, less the coincident.

### SiMPull threshold analysis

Single-channel estimations of aggregated species were carried out using the AF488 channel. A threshold intensity value was subtracted from all pixels of the image to exclude signal that is unlikely to originate from oligomeric species. This value was calculated as three standard deviations above the mean pixel signal of all non-coincident species identified via STAPull analysis. Particle detection and quantification was subsequently carried out using the ComDet v.0.5.5 plugin for ImageJ.

### Limit of detection analysis

100 fM to 10 nM α-syn aggregate concentration series were prepared in triplicate in 0.02 μm-filtered PBS (Fisher Scientific, 10209252) and the resultant aggregate numbers quantified by STAPull and SiMPull. For comparison to ThT fluorimetry, an additional 0.1-7 μM concentration series was prepared in 5 μM ThT-supplemented PBS and their fluorescence intensity measured using a Denovix DS-11 Fxt Fluorimeter with 470 nm excitation and 514-567 nm emission detection. Concentration calibration curves were generated using the mean detections and fit with a second order polynomial using OriginPro software (OriginLab). The limit of blank (LoB), which indicates the highest number of counts expected when no analyte is detected, was calculated as^13^:

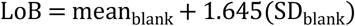

where the blank sample contained no protein. The limit of detection (LoD), which denotes the lowest analyte concentration likely to be reliably distinguished from the LoB, was subsequently calculated as:

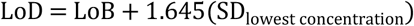

and the equivalent concentration derived from the standard curve equation. We report aggregate concentration based on the equivalent monomer starting concentration, which overestimates the oligomer fraction. The oligomer concentration was thus estimated as:

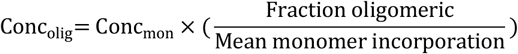

where fraction oligomeric is the proportion of total STAPull detections that were coincident, and mean monomer incorporation is the average coincident signal normalised by average non-coincident signal, both determined with a 5 nM α-syn aggregate sample.

### SAVE analysis

SAVE image analysis was performed using custom-written python code. Single-channel image series were mean intensity projected and a rolling background subtraction was applied. Local background was computed over an 11 × 11 kernel as the mean value less the global baseline as given by the intensity of the lowest 1% of pixels. The image was thresholded and binarized, using a threshold value equal to three standard deviations above the mean image signal when no protein or monomeric protein alone is present, and resulting features counted.

### Differentiation and culture of iPSC-derived mDA neurons

*SNCA* tripication iPS cell line, AST18, was derived from skin fibroblasts donated by a PD patient harbouring a triplication of a region on Chr4q22 encompassing the *SNCA* locus^31^. A full knock out of all four *SNCA* copies, AST18-7B, was derived from the parent triplication line via CRISPR^32^. iPS cell culture and directed differentation protocols used to generate mDA neuons were described previously by Chen et al 2019. Successful differentiation of mDA neurons was confirmed by probing cell type-specific identity markers (supplementary figure 6).

### CSF samples

CSF samples were collected from 7 patients with clinically-defined PD (aged 53-75, mean ± SD = 64 ± 7), 5 patients with clinically-defined AD (aged 43-68, mean ± SD = 60 ± 10) and 13 healthy individuals (aged 46-76, mean ± SD = 63 ± 7) based on internationally established criteria^33,34^. Samples were initially obtained for a previous biomarker study^22^. Full demographic, clinical and biomarker information is outlined in supplementary table 2. Sample collection and storage followed a standardized procedure (www.neurochem.gu.se/TheAlzAssQCProgram). Briefly, lumbar puncture was performed between 09:00-12:00. 15 mL of CSF were collected in sterile polypropylene tubes. The collected CSF samples were gently mixed to avoid gradient effects and centrifuged at 4,000 rpm for 10 min at 4 °C before dividing into 0.5 mL aliquots in Protein LoBind tubes, which were frozen on dry ice and stored at -80 °C. Blood-contaminated samples (>500 red blood cells [RBCs] per µL) were excluded. Time between sample collection, centrifugation and freezing was maximum 1 hour. The study was conducted in accordance with local clinical research regulations and with the provisions of the Helsinki declaration. Informed consent was obtained from all subjects, including access to their clinical data.

## Acknowledgements

We thank Kerry Tyson, Sarfaraj Topia and Hanna Hailu for reagent generation. The single-molecule instruments used in this study were funded by the UK Dementia Research Institute, UCB Biopharmaceuticals, and a kind donation from Dr. Jim Love. J.L. and R.S.S. were supported by UCB Biopharma S.P.R.L. C.L. was supported by an MRC National Productivity Investment Fund studentship. K.J. was funded by the BBSRCEastBIO doctoral training program (BB/M010996/1). A.C. was supported by the Engineering and Physical Sciences Research Council and the MRC through the Centre for Doctoral Training in Optical Medical Imaging (grant no. EP/L016559/1), the Rosetrees Trust and the Scottish Funding Council (H14052/SIRL ID: 691). SRB was supported by a Research Training Program Scholarship from the Australian Government (Department of Education, Skills and Employment). MS acknowledges funding from an Australian Research Council Discovery Project grant (DP200102463). S.G. is an MRC Senior Clinical Fellow (MR/T008199/1). T.K. was supported by MRC (MR/K017276/1) and Cure Parkinson’s.

## References

1. Chiti, F. & Dobson, C. M. Protein misfolding, functional amyloid, and human disease. Annu. Rev. Biochem. 75, 333–366 (2006).

2. Winner, B. et al. In vivo demonstration that alphasynuclein oligomers are toxic. Proc. Natl. Acad. Sci. U. S. A. 108, 4194–4199 (2011).

3. Lasagna-Reeves, C. A. et al. Tau oligomers impair memory and induce synaptic and mitochondrial dysfunction in wild-type mice. Mol. Neurodegener. 6, 39 (2011).

4. Tokuda, T. et al. Detection of elevated levels of α-synuclein oligomers in CSF from patients with Parkinson disease. Neurology 75, 1766–1772 (2010).

5. Walker, L. C. Proteopathic Strains and the Heterogeneity of Neurodegenerative Diseases. Annu. Rev. Genet. 50, 329–346 (2016).

6. Horrocks, M. H. et al. Single-Molecule Imaging of Individual Amyloid Protein Aggregates in Human Biofluids. ACS Chem. Neurosci. 7, 399–406 (2016).

7. Morten, M. J. et al. Quantitative super-resolution imaging of pathological aggregates reveals distinct toxicity profiles in different synucleinopathies. Proc. Natl. Acad. Sci. U. S. A. 119, e2205591119 (2022).

8. Lee, J.-E. et al. Mapping Surface Hydrophobicity of α-Synuclein Oligomers at the Nanoscale. Nano Lett. 18, 7494–7501 (2018).

9. Jain, A. et al. Probing cellular protein complexes using single-molecule pull-down. Nature 473, 484–488 (2011).

10. Je, G. et al. Endogenous Alpha-Synuclein Protein Analysis from Human Brain Tissues Using Single-Molecule Pull-Down Assay. Anal. Chem. 89, 13044–13048 (2017).

11. Chappard, A. et al. Single-molecule two-color coincidence detection of unlabeled alpha-synuclein aggregates. Angew. Chem. Int. Ed Engl. e202216771 (2023).

12. Hariri, A. A. et al. Improved immunoassay sensitivity and specificity using single-molecule colocalization. Nat. Commun. 13, 1–11 (2022).

13. Armbruster, D. A. & Pry, T. Limit of blank, limit of detection and limit of quantitation. Clin. Biochem. Rev. 29 Suppl 1, S49–52 (2008).

14. Chen, S. W. et al. Structural characterization of toxic oligomers that are kinetically trapped during α-synuclein fibril formation. Proceedings of the National Academy of Sciences 112, E1994–E2003 (2015).

15. van Steenoven, I. et al. α-Synuclein species as potential cerebrospinal fluid biomarkers for dementia with lewy bodies. Mov. Disord. 33, 1724–1733 (2018).

16. Majbour, N. K. et al. Oligomeric and phosphorylated alpha-synuclein as potential CSF biomarkers for Parkinson’s disease. Mol. Neurodegener. 11, 7 (2016).

17. Biancalana, M. & Koide, S. Molecular mechanism of Thioflavin-T binding to amyloid fibrils. Biochim. Biophys. Acta 1804, 1405–1412 (2010).

18. Lorenzen, N. et al. The Role of Stable α-Synuclein Oligomers in the Molecular Events Underlying Amyloid Formation. J. Am. Chem. Soc. 136, 3859–3868 (2014).

19. Singleton, A. B. et al. alpha-Synuclein locus triplication causes Parkinson’s disease. Science 302, 841 (2003).

20. Iljina, M. et al. Kinetic model of the aggregation of alpha-synuclein provides insights into prion-like spreading. Proc. Natl. Acad. Sci. U. S. A. 113, E1206–E1215 (2016).

21. Whiten, D. R. et al. Nanoscopic Characterisation of Individual Endogenous Protein Aggregates in Human Neuronal Cells. Chembiochem 19, 2033–2038 (2018).

22. Magdalinou, N. K. et al. A panel of nine cerebrospinal fluid biomarkers may identify patients with atypical parkinsonian syndromes. J. Neurol. Neurosurg. Psychiatry 86, 1240–1247 (2015).

23. Li, W. et al. Aggregation promoting C-terminal truncation of alpha-synuclein is a normal cellular process and is enhanced by the familial Parkinson’s disease-linked mutations. Proc. Natl. Acad. Sci. U. S. A. 102, 2162–2167 (2005).

24. Athauda, D. & Foltynie, T. Challenges in detecting disease modification in Parkinson’s disease clinical trials. Parkinsonism Relat. Disord. 32, 1–11 (2016).

25. Bousset, L. et al. Structural and functional characterization of two alpha-synuclein strains. Nature Communications 4, (2013).

26. Liu, P. et al. Quaternary Structure Defines a Large Class of Amyloid-β Oligomers Neutralized by Sequestration. Cell Rep. 11, 1760–1771 (2015).

27. Krejciova, Z. et al. Human stem cell-derived astrocytes replicate human prions in a PRNP genotype-dependent manner. J. Exp. Med. 214, 3481–3495 (2017).

28. Paleologou, K. E. et al. Phosphorylation at Ser-129 but not the phosphomimics S129E/D inhibits the fibrillation of alpha-synuclein. J. Biol. Chem. 283, 16895–16905 (2008).

29. Hoyer, W. et al. Dependence of alpha-synuclein aggregate morphology on solution conditions. J. Mol. Biol. 322, 383–393 (2002).

30. Reddy, B. S. & Chatterji, B. N. An FFT-based technique for translation, rotation, and scale-invariant image registration. IEEE Trans. Image Process. 5, 1266–1271 (1996).

31. Devine, M. J. et al. Parkinson’s disease induced pluripotent stem cells with triplication of the α-synuclein locus. Nat. Commun. 2, 440 (2011).

32. Chen, Y. et al. Engineering synucleinopathy-resistant human dopaminergic neurons by CRISPR-mediated deletion of the SNCA gene. Eur. J. Neurosci. 49, 510–524 (2019).

33. McKhann, G. et al. Clinical diagnosis of Alzheimer’s disease. Neurology 34, 939–939 (1984).

34. National Collaborating Centre for Chronic Conditions (UK). Parkinson’s Disease: National Clinical Guideline for Diagnosis and Management in Primary and Secondary Care. (Royal College of Physicians (UK)).

