## Supplementary Information for "Two-color coincidence single-molecule pull-down for the specific detection of disease-associated protein aggregates"

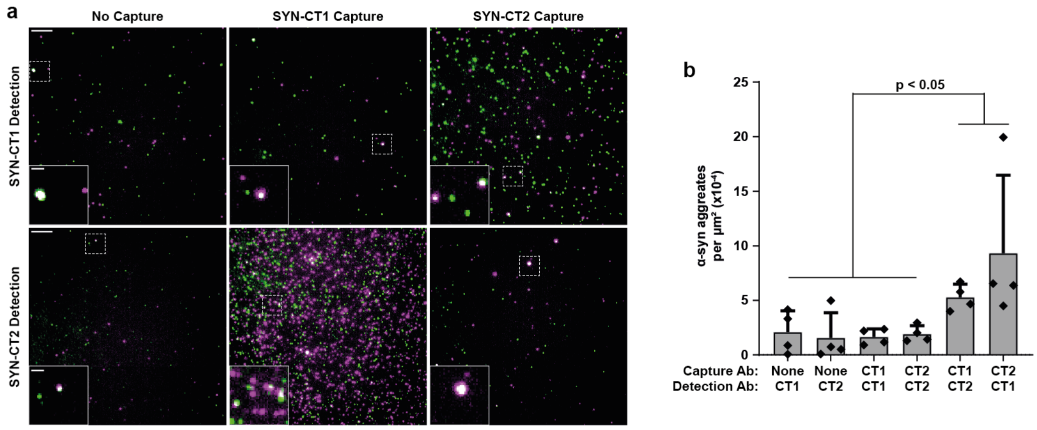

**Supplementary Figure 1. STAPull sensitivity is improved when different capture and detection antibodies are used. a** Representative fluorescence images of 10 nM α-syn subjected to conditions favouring aggregation for 48h, immobilized and visualized using different combinations of the SYN-CT1 and SYN-CT2 antibodies for capture and detection. **b** Quantification of colocalized events (mean ± SD, n=4, 16 FOV, total area = 64,119.71 $\mu m^2$). * $P < 0.05$ One-way ANOVA with Tukey multiple comparison test. (Note outlier in CT2 capture + CT1 detection sample was excluded for statistical analysis). Scale bars are 5 µm and 1 µm in length, for full view and inset images, respectively.

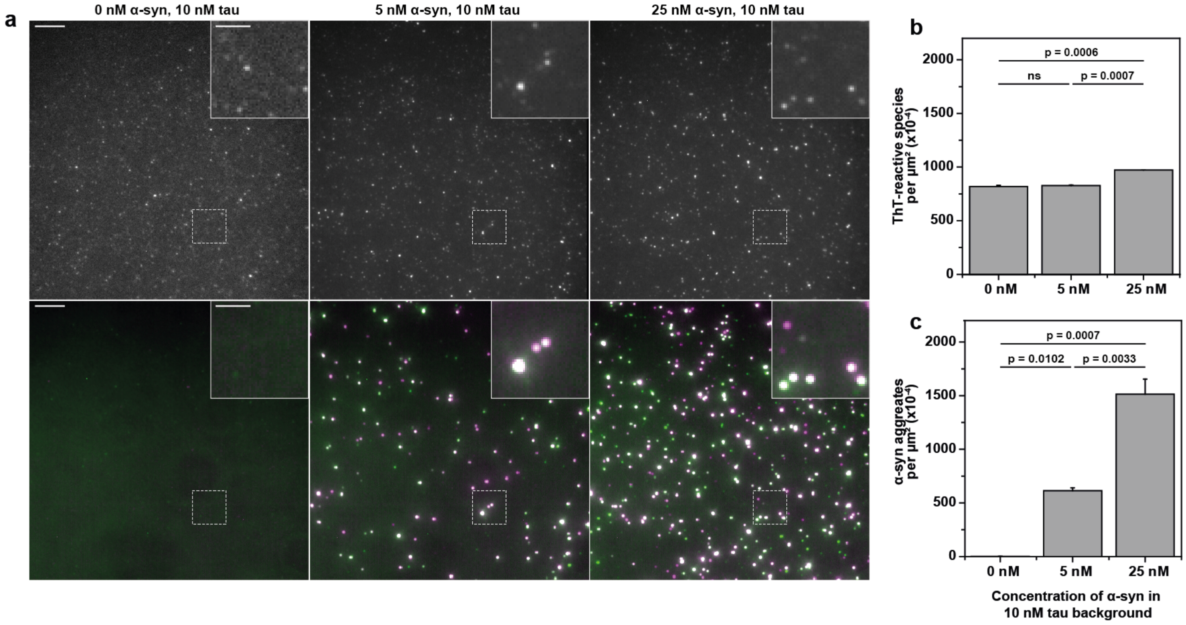

**Supplementary Figure 2. The STAPull detection antibody confers specificity. a** Representative TIRF microscopy images of mixed protein aggregate samples (10 nM tau with 0-25 nM α-syn, as indicated) pulled down by the SYN-CT1 anti-α-syn antibody. Detection used either SAVE ThT imaging to non-specifically detect all amyloid aggregates (top row) or STAPull with anti-α-syn SYN-CT2 to specifically detect α-syn aggregates (bottom row). Scale bars 5 µm in full-frame, 2 µm in inset. **b** Quantification of the number of ThT-reactive species and **c** STAPull coincident events per µm$^2$ for the dataset presented in a (mean ± SD, 64 technical repeats). Statistical analyses carried out using an unpaired student t-test.

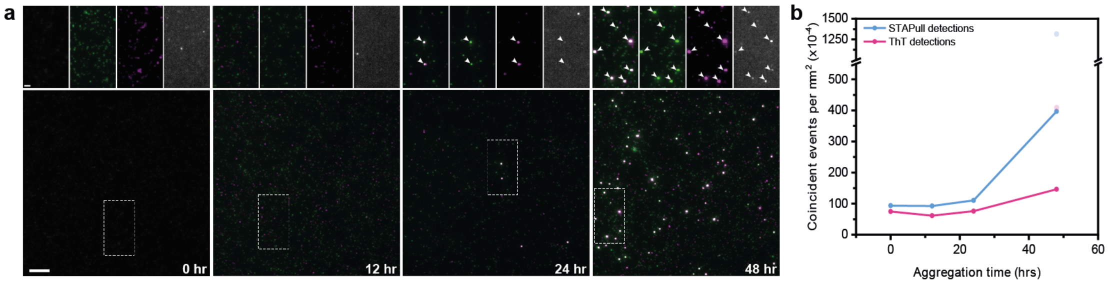

**Supplementary Figure 3. STAPull detects earlier aggregates than ThT. a** Representative STAPull images of α-syn incubated under aggregation-promoting conditions for 0-48 hours, as indicated. Magnified STAPull (left), single-channel (center) and ThT (right) counterstain images of the boxed region are shown above, coincident events representing aggregates are highlighted (white arrowhead) demonstrating a population of STAPull aggregates that are not ThT-reactive. Scale bars 5 µm in full-frame, 1 µm in inset. **b** Quantification of the mean number of aggregates per µm$^2$ over time as detected by STAPull (blue) or ThT (pink) for the dataset represented in a (mean ± SD, n = 3, 64 FOVs, outliers shown at 48 hours excluded from mean calculations).

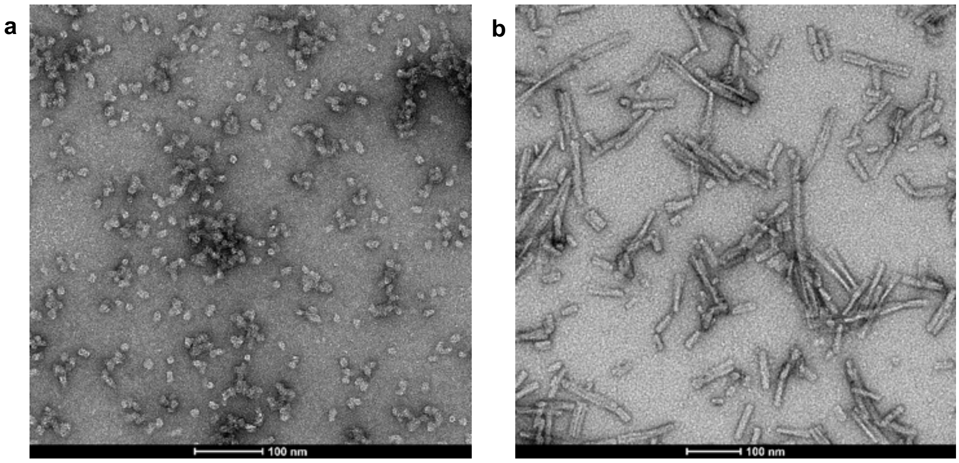

**Supplementary Figure 4. Structural validation of commercial α-syn constructs using transmission electron microscopy.** Representative transmission electron microscopy (TEM) images of commercially sourced recombinant **a** kinetically-trapped α-syn oligomers and **b** pre-formed fibrils. Samples were prepared using the 'direct application method' previously published[1]. Negative stain TEM images were acquired at 80 Kv on carbon coated 400 mesh copper grids using phosphotungstic acid and uranyl acetate stain.

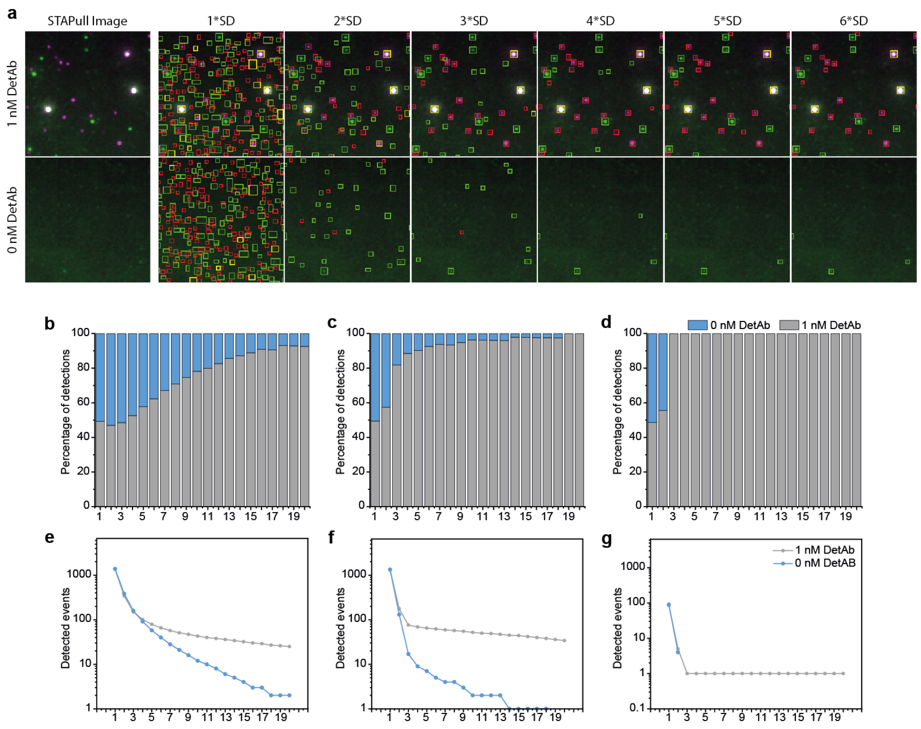

**Supplementary Figure 5. Empirical determination of the threshold value for ComDet particle detection. a** Representative STAPull images of 10 nM α-syn aggregates in the presence (top) or absence (bottom) of 1 nM SYN-CT2 detection antibody, alongside overlaid particle detections obtained for the same image using an intensity threshold set at 1-6 standard deviations above the mean, as indicated, with single channel (green or red boxes) and coincident (yellow boxes) detections shown. **b** The percent of combined detections that are specific to α-syn (grey, based on mean count in the presence of 1 nM detection antibody) and non-specific (blue, based on mean count in the absence of detection antibody) as a function of threshold value for AF488, **c** AF647, and **d** STAPull coincidence. **e-g** The mean particle detections for **e** AF488, **f** AF647, and **g** STAPull coincidence (over 64 technical repeats) in the presence (grey) or absence (blue) of 1 nM detection antibody.

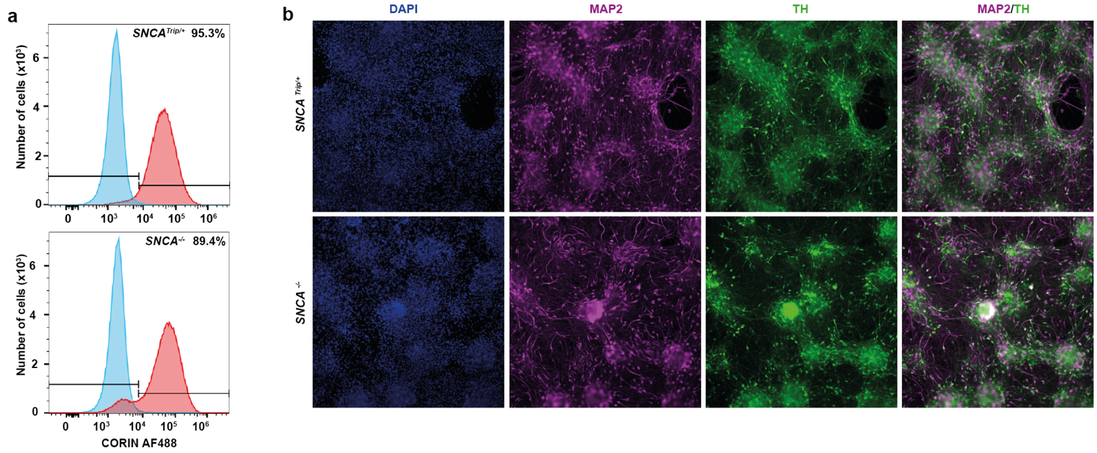

**Supplementary Figure 6. Differentiation of iPSCs into mDA neurons. a** Percentage of day 16 neural progenitor cells expressing cell surface protein CORIN, a floor plate identity marker for *SNCA*$^{Trip/+}$ (top) and *SNCA*$^{-/-}$ (bottom) cultures. **b** Immunostaining of day 98 mDA neurons differentiated from *SNCA*$^{Trip/+}$ (top) and *SNCA*$^{-/-}$ (bottom) iPSCs; DAPI (blue), Tyrosine Hydroxylase (green), Microtubule-associated protein 2 (Magenta). Scale bar is 200 µm in length.

| **PLL Surface** | **ANOVA: p = 1.54 x10-116** | | |
|---|---|---|---|
| Tukey matrix | Aβ | αSyn | Tau |
| Aβ | - | - | - |
| αSyn | <.00001 | - | - |
| Tau | 0.62459 | <.00001 | - |
| No Protein | 0.4375 | <.00001 | 0.99077 |
| **αSyn pull-down** | **ANOVA: p = 1.69 x10-213** | | |
| Tukey matrix | Aβ | αSyn | Tau |
| Aβ | - | - | - |
| αSyn | <.00001 | - | - |
| Tau | 0.99991 | <.00001 | - |
| No Protein | 0.99757 | <.00001 | 0.99926 |
| **Tau pull-down** | **ANOVA: p = 1.65 x10-167** | | |
| Tukey matrix | Aβ | αSyn | Tau |
| Aβ | - | - | - |
| αSyn | 0.99639 | - | - |
| Tau | <.00001 | <.00001 | - |
| No Protein | 0.99208 | 0.9999 | <.00001 |

**Supplementary Table 1. ANOVA post hoc statistical significance probabilities of STAPull specificity.** Matrices detail the Tukey probability of significant differences between the mean density of aggregates detected on a PLL-coated surface (top), an α-syn-specific STAPull surface (middle), or a tau-specific STAPull surface when exposed to different protein aggregates (refer to figure 3).

| Clinical data | | | | | ELISA | | | | | SAVE | STAPull | |
|---|---|---|---|---|---|---|---|---|---|---|---|---|
| Diagnosis | Age | Disease duration | HY Score | MMSE | t-protein | t-tau | p-tau | Aβ | **α**-syn | ThT species per **μ**$m^2$ | Total **α**-syn (fold above baseline) | **α**-syn aggregates (fold above baseline) |
| Healthy | 69 | | | | 0.7 | >1,200 | 100 | 726 | 3144 | 0.0016 | 0.788 | 6.054 |
| Healthy | 69 | | | | 0.53 | 818 | 75 | 1260 | 3472 | 0.0022 | 0.859 | 6.433 |
| Healthy | 67 | | | | 0.3 | 335 | 100 | 338 | 1830 | 0.0049 | 0.749 | 4.114 |
| Healthy | 64 | | | | 0.29 | 230 | 41 | 843 | 1154 | 0.0069 | 0.681 | 4.725 |
| Healthy | 64 | | | | 0.31 | 185 | 28 | 771 | 1413 | 0.0055 | 0.449 | 4.215 |
| Healthy | 63 | | | | 0.28 | 189 | 29 | 1148 | 1178 | 0.0045 | 1.698 | 3.207 |
| Healthy | 61 | | | | 0.22 | 241 | 32 | 865 | 1535 | 0.0174 | 0.807 | 3.694 |
| Healthy | 76 | | | | 0.65 | 306 | 33 | 305 | 1206 | 0.0058 | 0.868 | 3.964 |
| Healthy | 62 | | | | 0.25 | 499 | 54 | 918 | 3071 | 0.0038 | 1.722 | 6.361 |
| Healthy | 59 | | | | 0.49 | 107 | 20 | 677 | 1015 | | 1.045 | 4.263 |
| Healthy | 64 | | | | 0.46 | 402 | 40 | 1199 | 2189 | 0.0046 | 0.990 | 4.656 |
| Healthy | 46 | | | | 0.85 | 186 | 31 | 521 | 1407 | 0.0026 | 0.602 | 4.157 |
| Healthy | 61 | | | | 0.3 | 301 | 44 | 1345 | 1796 | 0.0043 | 0.443 | 1.838 |
| AD | 64 | 3 | | | 0.58 | 417 | 40 | 727 | 1238 | | 0.797 | 5.856 |
| AD | 62 | 2 | | | 0.52 | 329 | 45 | 863 | 2192 | | 0.711 | 4.813 |
| AD | 61 | 1 | | | 0.45 | 1092 | 128 | 445 | 2486 | | 0.809 | 4.365 |
| AD | 68 | 1 | | | 0.8 | 742 | 57 | 379 | 1400 | | 0.989 | 5.176 |
| AD (PSEN1) | 43 | 2 | | | 0.27 | >1,200 | 123 | 385 | 1786 | | 0.774 | 4.706 |
| iPD | 68 | 20 | 4 | 30 | 0.55 | 247 | 33 | 747 | 1763 | | 1.035 | 8.868 |
| iPD | 75 | 14 | 4 | 30 | 1.07 | 380 | 49 | 1157 | 2306 | 0.0148 | 2.453 | 7.683 |
| iPD | 58 | 6 | 2 | 30 | 0.34 | 471 | 50 | 770 | 1781 | 0.0129 | 1.523 | 6.422 |
| iPD | 69 | 11 | 3 | 30 | 0.29 | 516 | 39 | 1345 | 2918 | 0.01 | 1.879 | 6.743 |
| iPD | 53 | 5 | 1 | 28 | 0.46 | 322 | 30 | 921 | 1384 | 0.0237 | 2.323 | 7.672 |
| iPD | 60 | 6 | 2 | 30 | 0.44 | 135 | 18 | 744 | 777 | 0.0148 | 3.123 | 8.318 |
| iPD | 65 | 15 | 3 | 27 | 0.37 | 509 | 62 | 732 | 2311 | 0.0177 | 1.792 | 10.672 |

**Supplementary Table 2. Clinical details of patients and controls along with CSF biomarker analysis.** The CSF samples used in this study were collected for a previous biomarker study[2], and were obtained from UCL Queen Square Institute of Neurology. Informed consent was obtained from all subjects, including access to their clinical data. No additional ethics approval was required for our use of the CSF after obtaining it from the UCL Queen Square Institute of Neurology. The PD patients were classified according to their HY (Hoehn & Yahr) grade. The total protein, total tau, phosphorylated tau, Aβ and α-syn were measured by ELISA[2]. The SAVE counts, where available, are from Horrocks et al[3].

| Antibody | Target | Human α-syn monomer $k_D$ [nM] | Human α-syn fibril $k_D$ [pM] |
|---|---|---|---|
| SYN-NT1 | α-syn N-terminal | 41.0 | 463 |
| SYN-CT1 | α-syn early C-terminal | 0.18 | 40 |
| SYN-CT2 | α-syn late C-terminal | 10.0 | 30 |
| Anti-tau | Tau | n/a | n/a |

**Supplementary Table 3. Antibodies used in this study.** Affinities to monomer and fibrillar α-syn were measured by SPR.

### Supplementary Information References

1. Doane, F. W. & Anderson, N. *Diagnostic Virology—A Practical Guide And Atlas*. (Cambridge University Press, 1997).

2. Magdalinou, N. K. *et al.* A panel of nine cerebrospinal fluid biomarkers may identify patients with atypical parkinsonian syndromes. *J. Neurol. Neurosurg. Psychiatry* **86**, 1240–1247 (2015).

3. Horrocks, M. H. *et al.* Single-Molecule Imaging of Individual Amyloid Protein Aggregates in Human Biofluids. *ACS Chem. Neurosci.* **7**, 399–406 (2016).
